# Mutagenesis of the Ammonium Transporter *AcAmt* Reveals a Reproductive Role and a Novel Ammonia-Sensing Mechanism in the Malaria Vector Mosquito *Anopheles coluzzii*

**DOI:** 10.1101/2021.01.29.428879

**Authors:** Zi Ye, Feng Liu, Stephen T. Ferguson, Adam Baker, R. Jason Pitts, Laurence J. Zwiebel

**Affiliations:** Department of Biological Sciences, Vanderbilt University, Nashville, TN 37235, USA; Department of Biology, Baylor University, Waco, TX 76706, USA

**Keywords:** Olfaction, Ammonium transporter, *Anopheles coluzzii*, CRISPR/Cas9, Mosquito reproduction, Coeloconic sensilla

## Abstract

Anopheline mosquitoes are the sole vectors of malaria and rely on olfactory cues for host seeking in which ammonia derived from human sweat plays an essential role. To investigate the function of the *Anopheles coluzzii* ammonium transporter (*AcAmt*) in the mosquito olfactory system, we generated an *AcAmt* null mutant line using CRISPR/Cas9. *AcAmt*^-/-^ mutants displayed a series of novel phenotypes compared with wild-type mosquitoes including significantly lower insemination rates during mating and increased mortality during eclosion. Furthermore, *AcAmt*^-/-^ males showed significantly lower sugar consumption while *AcAmt*^-/-^ females and pupae displayed significantly higher ammonia levels than their wild-type counterparts. Surprisingly, in contrast to previous studies in *Drosophila* that revealed that the mutation of the ammonium transporter (*DmAmt*) induces a dramatic reduction of ammonia responses in antennal coeloconic sensilla, no significant differences were observed across a range of peripheral sensory neuron responses to ammonia and other odorants between wild-type and *AcAmt*^-/-^ females. Taken together, these data support the existence of a unique ammonia-sensing mechanism in mosquitoes and that the ammonium transporter may be an important molecular target for vector control.

**Key Messages:**

- Mutagenesis of *An. coluzzii* ammonium transporter *AcAmt* followed by comprehensive electrophysiological investigation suggest a novel ammonia-sensing pathway in *Anopheles* mosquitoes.
- *AcAmt*^-/-^ mutants displayed significant deficiencies in reproduction and eclosion, which are likely due to elevated ammonia levels and reduced ability of sugar feeding.
- *An. coluzzii* coeloconic sensilla primarily detect amines and acids.

## Introduction

Several species of Anopheline mosquitoes make up the primary vectors of *Plasmodium* parasites that are the causative agents for human malaria resulting in hundreds of thousands of deaths worldwide every year (World Health Organization 2019). Pathogen transmission occurs exclusively as a consequence of the blood meals that female mosquitoes require in order to complete their reproductive cycles. The mosquito’s olfactory system provides the ability to sense and discriminate a broad spectrum of semiochemical cues that drive host preference and seeking behaviors that ultimately lead to blood feeding (Zwiebel and Takken 2004; Carey and Carlson 2011; Montell and Zwiebel 2016). In that process, ammonia along with several carboxylic acids derived from human sweat act as attractants that promote mosquito-human interactions (Smallegange et al. 2011). Anopheline females are attracted to ammonia without the presence of other sweat-derived cues (Braks et al. 2001).

A complex array of molecular components, which most notably include two classes of chemosensory receptors, odorant receptors (ORs) and ionotropic receptors (IRs), are highly expressed on the antennae and other olfactory appendages of Anopheline females where they have been implicated in the neuronal sensitivity to a range of odorant stimuli (Pitts et al. 2004; Pitts et al. 2017; Sun et al. 2020). While *Ir92a* has been characterized in *Drosophila* as an ammonia receptor expressed in antennal neurons, the molecular pathway of ammonia detection in mosquitoes has remained cryptic due to the lack of a direct homolog to *Drosophila Ir92a* (Benton et al. 2009; Min et al. 2013).

The transportation of ammonium in bacteria, insects, and other animals occurs through the aptly named ammonium transporter (Amt) (Andrade and Einsle 2007; Tremblay and Hallenbeck 2009; Pitts et al. 2014a). While bacteria rely on Amt for both ammonium uptake and diffusion (Thomas et al. 2000; Soupene et al. 2002), in *Aedes aegypti, AeAmt1* expressed in the anal papillae is involved in ammonium excretion and *AeAmt1* RNAi treated larvae display significantly higher concentrations of ammonium ions in the hemolymph than wild-type mosquitoes (Chasiotis et al. 2016; Durant and Donini 2018). More recently, several studies focused on Amt revealed a novel function in mediating ammonia sensitivity in insect chemosensory systems (Menuz et al. 2014; Pitts et al. 2014a; Delventhal et al. 2017). In *Drosophila melanogaster*, the ammonium transporter (*DmAmt*) is expressed in the auxiliary cells of coeloconic ac1 sensilla, in which null mutations result in a loss of antennal sensitivity to ammonia (Menuz et al. 2014). Furthermore, the expression of *DmAmt* in auxiliary cells, as opposed to the olfactory sensory neurons (OSNs), suggested it may not be a molecular sensor of ammonia but rather could be involved in ammonium clearance which prevents neuron desensitization. Studies in *Anopheles coluzzii* (formerly *An. gambiae;* Coetzee et al. 2013) suggested that AcAmt facilitates cross-membrane transport of ammonium ions in a heterogeneous expression system (Pitts et al. 2014a), and, importantly, *AcAmt* is localized in the antennal auxiliary cells of basiconic (grooved pegs) and coeloconic sensilla (Ye et al. 2020). These data suggest there may be conserved functionality between *Drosophila* and mosquitoes, in the latter case where ammonia sensing pathways plays a substantial role in host seeking (Ye et al. 2020). Thus far, technical difficulties in gene editing in *Anopheles* mosquitoes has precluded elucidation of the olfactory function of AcAmt *in vivo*.

Here we used CRISPR/Cas9 to generate an *AcAmt* null mutant line to examine the hypothesis that *AcAmt* is essential for ammonia responses in *Anopheles* mosquitoes. Surprisingly, *AcAmt* mutants failed to display a significant difference in ammonia peripheral responses in antennal single sensillum recordings (SSRs) as well as in electroantennogram (EAG) and electrolabellogram (ELG) assays compared with wild-type *An. coluzzii*. These results suggest a divergence of ammonia-sensing pathways between *Drosophila* and mosquitoes. Furthermore, we observed *AcAmt* null mutants to be dramatically less efficient in mating and pupal eclosion. A series of behavioral and biochemical assessments were undertaken to investigate the potential mechanisms underlying these behavioral defects.

## Material and Methods

### Mosquito rearing

*An. coluzzii* (SUA 2La/2La), previously known as *Anopheles gambiae sensu stricto* “M-form”(Coetzee et al. 2013), originated from Suakoko, Liberia, were reared using previously described protocols (Fox et al., 2001; Qiu et al., 2004). Briefly, all mosquito lines were reared at 27°C, 75% relative humidity under a 12:12 light:dark cycle (11 h ~250 lux full light, 11 h darkness, with 1 h dawn/dusk gradient transitions in between) and supplied with 10% sugar water in the Vanderbilt University Insectary (Fox et al. 2001; Suh et al. 2016). Mosquito larvae were reared in 500mL distilled water with 100 larvae per rearing pan. Larval food was prepared by dissolving 0.12g/mL Kaytee Koi’s Choice premium fish food (Chilton, WI, US) and 0.06g/mL yeast in distilled water and incubating at 4°C overnight for fermentation. For 0-to 4-day-old larvae, 0.12mL of larval food solution was added daily into each rearing pan; for larvae ≥5 days old, 0.16mL was added.

### Mosquito mutagenesis

CRISPR/Cas9 gene editing in *An. coluzzii* was carried out as previously described (Liu et al. 2020), with minor modifications. The CRISPR gene-targeting vector was a kind gift from Dr. Andrea Crisanti of Imperial College London, UK (Hammond et al. 2016). The single guide RNA (sgRNA) sequences for *AcAmt* gene *Exon 1* and *Exon 4* were designed by CHOPCHOP (http://chopchop.cbu.uib.no/) with high efficiency (**Supplementary Table 1**) and were synthesized (Integrated DNA Technologies, Coralville, IA) and subcloned into the CRISPR vector by Golden Gate cloning (New England Biolabs, Ipswich, MA). The homologous templates were constructed based on a pHD-DsRed vector (a gift from Kate O’Connor-Giles; Addgene plasmid #51434; http://n2t.net/addgene:51434; RRID:Addgene 51434), in which the 2-kb homologous arms extending either direction from the double-stranded break (DSB) sites were PCR amplified (**Supplementary Table 1**) and sequentially inserted into the *AarI* and *SapI* restriction sites on the vector, respectively.

The microinjection protocol was carried out as described (Pondeville et al. 2014; Ye et al. 2020). Briefly, newly laid (approximately 1-h old) embryos of the wild-type *An. coluzzii* were immediately collected and aligned on a filter paper moistened with 25mM sodium chloride solution. All the embryos were fixed on a coverslip with double-sided tape, and a drop of halocarbon oil 27 (Sigma-Aldrich, St. Louis, MO) was applied to cover the embryos. The coverslip was further fixed on a slide under a Zeiss Axiovert 35 microscope with a 40X objective (Zeiss, Oberkochen, Germany). The microinjection was performed using Eppendorf FemtoJet 5247 and quartz needles (Sutter Instrument, Novato, CA). The gene targeting vectors at 300ng/μL were co-injected with the homologous template at 300ng/μL. The injected embryos were placed into deionized water with artificial sea salt (0.3g/L) and reared under lab conditions.

First-generation (G0) injected adults were separated based on sex and crossed with 5X wild-type sex counterparts. Their offspring (F1) were screened for DsRed-derived red eye fluorescence. Red-eyed F1 males were individually crossed with 5X wild-type females to establish a stable mutant line. PCR analyses of all individuals were performed (after mating) to validate the fluorescence marker insertion using primers that cover the DSB site (**Supplementary Table 1**). The PCR products were further sequenced to confirm the accurate insertion. The heterozygous mutant lines were back-crossed with the wild-type partners for at least eight generations before putative homozygous individuals were manually screened for DsRed-derived red-eye fluorescence intensity. Putative homozygous mutant individuals were mated to each other before being sacrificed for genomic DNA extraction and PCR analyses (as above) to confirm their genotypes.

### Single sensillum recording (SSR)

SSR was carried out as previously described (Liu et al. 2013) with minor modifications. Non-blood-fed female mosquitoes (4-10 days post-eclosion) were mounted on a microscope slide (76 x 26 mm) (Ghaninia et al. 2007). The antennae were fixed using double-sided tape to a cover slip resting on a small bead of dental wax to facilitate manipulation, and the cover slip was placed at approximately 30 degrees to the mosquito head. Once mounted, the specimen was placed under an Olympus BX51WI microscope and the antennae viewed at high magnification (1000X). Two tungsten microelectrodes were sharpened in 10% KNO_2_ at 10 V. The grounded reference electrode was inserted into the compound eye of the mosquito using a WPI micromanipulator, and the recording electrode was connected to the preamplifier (10X, Syntech) and inserted into the shaft of the olfactory sensillum to complete the electrical circuit to extracellularly record OSN potentials (Den Otter et al. 1980). Controlled manipulation of the recording electrode was performed using a Burleigh micromanipulator (Model PCS6000). The preamplifier was connected to an analog-to-digital signal converter (IDAC-4, Syntech), which in turn was connected to a computer for signal recording and visualization.

Stock odorants of highest available purity were diluted in paraffin oil to make 10^-2^ (v/v) working solutions. Ammonium hydroxide (Sigma-Aldrich, St. Louis, MO) was serially diluted in water to 0.01, 0.05, 0.1, 0.5, 1, and 5% ammonia solutions. For each odorant, a 10-μL aliquot was applied onto a filter paper (3 x 50mm), which was then inserted into a Pasteur pipette to create the stimulus cartridge. A sample containing the solvent (water/paraffin oil) alone served as the control. The airflow across the antennae was maintained at a constant 20 mL/s throughout the experiment. Purified and humidified air was delivered to the preparation through a glass tube (10-mm inner diameter) perforated by a small hole 10cm away from the end of the tube into which the tip of the Pasteur pipette could be inserted. The stimulus was delivered to the sensilla by inserting the tip of the stimulus cartridge into this hole and diverting a portion of the air stream (0.5L/min) to flow through the stimulus cartridge for 500ms using a stimulus controller (Syntech). The distance between the end of the glass tube and the antennae was ≤ 1cm. Signals were recorded for 10s starting 1s before stimulation, and the action potentials were counted off-line over a 500-ms period before and after stimulation. Spike rates observed during the 500-ms stimulation were subtracted from the spontaneous activities observed in the preceding 500ms and counts recorded in units of spikes/sec.

### Electroantennogram (EAG) and Electrolabellogram (ELG)

The EAG and ELG protocols were derived from previous studies (Kwon et al. 2006; Suh et al. 2016; Sun et al. 2020). Briefly, a non-blood-fed, 5-to 10-day-old female mosquito was decapitated with forceps. Two sharp borosilicate glass (1B100F-3; World Precision Instruments, Sarasota, FL) electrodes were prepared using an electrode puller (P-2000; Sutter Instruments, Novato, CA) and filled with Ringer solution (96mM NaCl, 2mM KCl, 1mM MgCl_2_, 1mM CaCl_2_, 5mM HEPES, pH = 7.5), in which a AgCl-coated sliver wire was placed in contact to complete a circuit with a reference electrode inserted into the back of the head. Antennal/labellar preparations were continuously exposed to a humidified air flow (1.84L/min) transferred through a borosilicate glass tube (inner diameter = 0.8cm) that was exposed to the preparation at a distance of 10mm. Stimulus cartridges were prepared by transferring 10μl of test or control stimuli solutions to filter paper (3 x 50mm), which was then placed inside a 6-inch Pasteur pipette. Odorant stimuli were delivered to antennal preparations for 500ms through a hole placed on the side of the glass tube located 10cm from the open end of the delivery tube (1.08L/min), where it was mixed with the continuous air flow using a dedicated stimulus controller (Syntech, Hilversum, The Netherlands). An air flow (0.76L/min) was simultaneously delivered from another valve through a blank pipette into the glass tube at the same distance from the preparation in order to minimize changes in flow rate during odor stimulation. The resulting signals were amplified 10x and imported into a PC via an intelligent data acquisition controller (IDAC-232; Syntech, Hilversum, The Netherlands) interface box, and the recordings were analyzed offline using EAG software (EAG Version 2.7, Syntech, Hilversum, The Netherlands). Maximal response amplitudes of each test stimuli were normalized after dividing by the control (solvent alone) responses.

### Pupation and eclosion rate quantification

Each replicate consisted of 80-100 newly hatched 1^st^ instar larvae reared under the same conditions with a density of 10 larvae/50μL dH_2_O. Pupae from each replicate were then collected into a mosquito bucket and allowed to eclose. Total pupae were counted and divided by the initial 1^st^ instar larval counts to calculate the pupation rate. The successfully eclosed adults were counted and divided by the pupal counts to measure the eclosion success rate.

### Mating bioassay

Newly emerged wild-type females and males were separated for 1 day. 15 females and 10 males were then placed in a rearing bucket and allowed to freely mate for 5 days. All surviving females were then collected and their spermathecae were dissected under a compound microscope. The spermathecae were then placed in the buffer (145mM NaCl, 4mM KCl, 1mM MgCl_2_, 1.3mM CaCl_2_, 5mM D-glucose, 10mM HEPES) (Pitts et al. 2014b) with 300nM DAPI and a cover slip was used to gently press and break the spermathecae to release the sperm. The spermathecae were examined to assess the insemination status under a 1000X compound microscope (BX60; Olympus, Tokyo, Japan). The insemination rate was calculated by dividing the number of inseminated females by the total number of females in each bucket.

### Mosquito locomotor activity bioassay

Individual adult mosquitoes (3-to 9-days old) were first anesthetized on ice, then placed in wells of a six-well CytoOne tissue-culture plate (CC7672-7506; USA Scientific, Ocala, FL), and thereafter allowed to recover for at least 30 min prior to trial start. The wells were supplied with a cotton ball soaked in 0.5mL of 10% sugar water. Activity was digitally recorded and analyzed starting at ZT12 (the onset of the dark cycle) and continued through to ZT17. Activity recordings were collected with VideoVelocity software (v3.7.2090, Vancouver, Canada) at one image per second using a USB camera (Spinel, Newport Beach, CA) with built-in 850nm IR light placed ~20cm above the six-well plate.

Digital recordings were analyzed *post hoc* using EthoVision software (v8.5, Noldus, Wageningen, NL) to generate the following activity/mobility parameters: (1) distance travelled, defined as movement of the center-point of the animal (cm); (2) time spent moving relative to time spent not moving using the following parameters defined according to the software: averaging interval, 1 sample; start velocity, 1.00cm/s; stop velocity, 0.90cm/s; (3) clockwise and counterclockwise turns, defined as a cumulative turn angle of 180° with a minimum distance travelled by the animal of at least 0.5cm, with turns in the opposite direction of less than 45.00° ignored; and (4) time the mosquito spent in the half of the well containing the sugar water.

### Capillary feeder (CAFE) bioassay

The CAFE bioassay was conducted following a previous study with minor modifications (Dennis et al. 2019). Each trial started at ZT12 and ended at ZT18 for 6h. Four 4-to 8-day-old mosquitoes were provided with water but otherwise fasted for 22h before being anesthetized on ice briefly and placed into a *Drosophila* vial (24.5mm x 95mm; Fisher Scientific, Waltham, MA). A borosilicate glass capillary (1B100F-3; World Precision Instruments, Sarasota, FL) was filled with 10% sucrose water and embedded into a cotton plug. The vial opening was then blocked with the cotton plug and the capillary was placed slightly protruding from the plug into the vial for mosquitoes to feed on. The sugar level in the capillary was compared before and after each trial to generate the initial sugar consumption value. At least four control vials with no mosquitoes inside were used to assess the evaporation at the same time. The final sugar consumption was calculated by subtracting the evaporation from the initial sugar consumption value.

### Mass measurements

Individual 3-to 6-day-old mosquitoes were briefly anesthetized on ice and weighed using a XSR Analytical Balance (Mettler Toledo, Columbus, OH).

### Ammonia quantifications

The total ammonia context of adult and pupal stage *An. coluzzii* was assessed according to (Scaraffia et al. 2005) with minor modifications. Here, two 3-to 5-day-old adults or a single ≥1-day-old pupa were homogenized in 150μL distilled water and centrifuged at max speed in a table centrifuge for 2min at 4°C. 100μL supernatant was used for ammonia level measurement following the manufacturer’s instructions of the Ammonia Reagent Set (Pointe Scientific, Canton, MI) (Scaraffia et al. 2005).The absorbance was read at 340nm wavelength using a SmartSpec 3000 spectrophotometer (Bio-Rad, Hercules, CA) and compared with an ammonia standard curve prepared with ammonium chloride to calculate the ammonia concentration.

### Carbohydrate quantification

The total carbohydrate content of adult and pupal stage *An. coluzzii* was assessed according to (Ahmed 2013; Ellison et al. 2015) with minor modifications. Here, four 3-to 6-day-old mosquitoes or ≥1-day-old pupae were collected between ZT11 and ZT12 and homogenized in 200μL ddH_2_O; the homogenate was centrifuged at maximum speed for 1min at 4°C. 10μL of the supernatant was collected from the homogenate and added to a phenol solution of 195μL ddH_2_O and 5μL 100% phenol. 500μL sulfuric acid was subsequently added to the solution and briefly vortexed. The colorimetric reaction stood at room temperature for 10min and then the absorbance was read at 490nm wavelength using a SmartSpec 3000 spectrophotometer (Bio-Rad, Hercules, CA). The absorbance was compared with a standard curve prepared with glucose to calculate the carbohydrate content.

## Results

### Generation of the *AcAmt* null mutant

A complete *AcAmt* null mutant strain was generated using CRISPR/Cas9 gene editing via embryonic microinjection of two targeting plasmids expressing Cas9 and dual sgRNAs along with a homology template to knock-in a *3xP3-DsRed* eye-specific red fluorescence marker between two *AcAmt* DSB sites (Liu et al. 2020). The two sgRNAs targeted sequences at the start of both *Exon 1* and *Exon 4* to remove the majority of the three exons in between the DSBs of the *AcAmt* coding region (**Supplementary Table 1**). The 2kb homology arms were designed to extend outward from the two DSB sites to insert the *3xP3-DsRed* fluorescence marker (**Figure 1A**). The successful knock-out/knock-in was molecularly confirmed in progeny using both PCR (**Figure 1B**) and DNA sequencing. Homozygous and heterozygous individuals from subsequent backcross generations were selected based on the intensity of red fluorescence that directly correlates to the copy number of *3xP3-DsRed* alleles.

**Figure 1.**
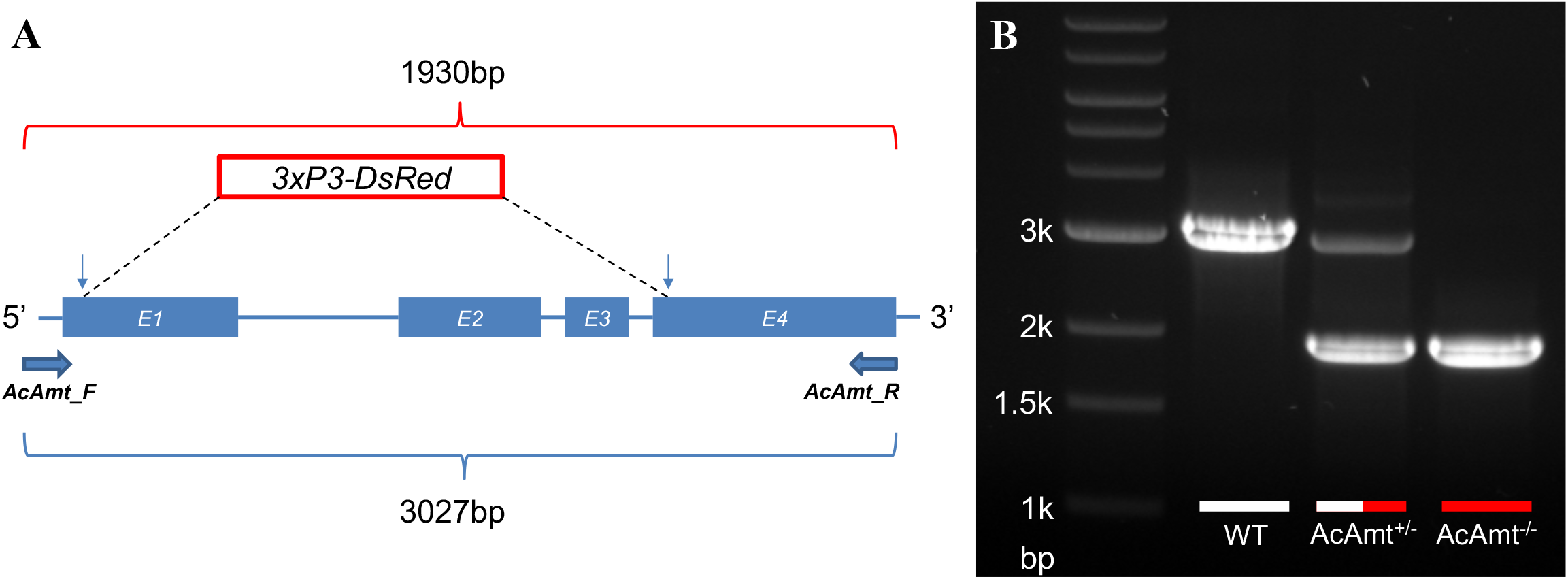
(**A**) Schematics of the CRISPR/Cas9 strategy that induced two double-stranded breaks (indicated by vertical arrows) on *Exon 1* (*E1*) and *Exon 4* (*E4*). A homology template was introduced to replace the sequences in between two double-stranded break sites with a red fluorescence marker *3xP3-DsRed*. A pair of primers (*AcAmt_F* and *AcAmt_R*) were used to determine the successful genetic manipulation. In theory, the wild-type produces a 3027-bp amplicon whereas the mutant renders a 1930-bp amplicon; (**B**) PCR determination of CRISPR/Cas9-mediated mutagenesis in the wild-type (WT), the heterozygotes (*AcAmt*^+/-^), and the homozygotes *AcAmt*^-/-^).

### Olfactory responses to ammonia

*AcAmt* expression has been localized to ammonia-sensitive antennal coeloconic sensilla and grooved pegs (Ye et al. 2020), which corresponds to the ammonia-sensing deficit in ac1 sensilla in *Drosophila* (Menuz et al. 2014). Here, SSR studies were carried out to examine whether responses to ammonia in these sensilla are affected by the *AcAmt*^-/-^ mutation (**Figure 2A&2B**). Surprisingly, and in contrast to the significant electrophysiological deficits observed in *DmAmt*^-/-^ mutants (Menuz et al. 2014), indistinguishable dose-dependent responses to ammonia were observed in coeloconic sensilla (**Figure 2C**) and grooved pegs (**Figure 2D**) in both wild-type and *AcAmt*^-/-^ females. Sensillar responses to repeated stimulations of ammonia were also assessed in order to saturate the sensillar lymph and potentially uncover a requirement for the putative clearance function of Amt (Menuz et al. 2014). Despite this additional challenge, no significant differences were observed in SSR responses across coeloconic sensilla (**Figure 2E**) and grooved pegs (**Figure 2F**) in wild-type and *AcAmt*^-/-^ female antennae. To investigate whether the *AcAmt*^-/-^ mutation alters sensillar responses to other odorants, we characterized the response profiles of coeloconic sensilla to an odorant panel of amines, acids, ketones, aldehydes, and alcohols (**Figure 3A**). Most amines and acids evoked strong, albeit not significantly different, responses in wild-type and *AcAmt*^-/-^ females (**Figure 3B**), as opposed to the weak responses elicited by other general odorants (**Figure 3C**).

**Figure 2.**
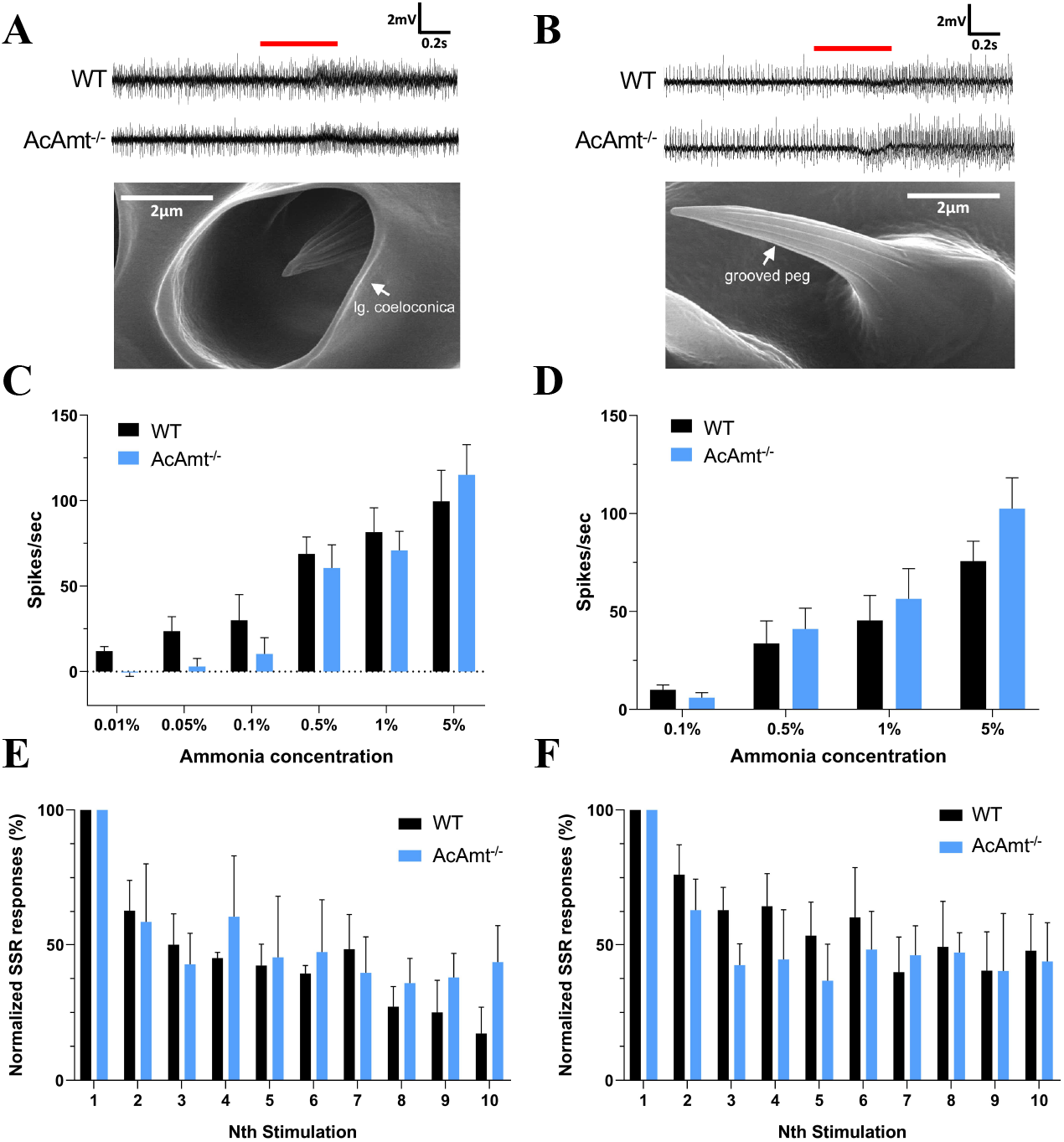
(**A**) Representative single-sensillum recordings from representative single-sensillum recording responses of coeloconic sensilla to 1% ammonia. Red bar indicates the duration of stimulations (0.5s). The scanning electron microscopy image showing the structure of a coeloconic sensillum is adopted from (Pitts and Zwiebel 2006); (**B**) Representative single-sensillum recording responses of grooved pegs to 0.5% ammonia. Red bar indicates the duration of stimulations (0.5s). The scanning electron microscopy image showing the structure of a grooved peg is adopted from (Pitts and Zwiebel 2006); (**C**) Single-sensillum responses of coeloconic sensilla to ammonia at different concentrations (N=5-7 for each concentration); (**D**) Single-sensillum responses of grooved pegs to ammonia at different concentrations (N=7-9 for each concentration); (**E**) Multiple single-sensillum responses of coeloconic sensilla to 1% ammonia with 5-s intervals (N=3). The responses were normalized to the fraction of the first stimulation; (**F**) Multiple single-sensillum responses of grooved pegs to 0.5% ammonia with 5-s intervals (N=3-5). The responses were normalized to the fraction of the first stimulation. Multiple t-tests with Holm-Sidak method suggest no significant differences (P > 0.05) between the wild-type and *AcAmt*^-/-^ Error bars = Standard error of the mean.

**Figure 3.**
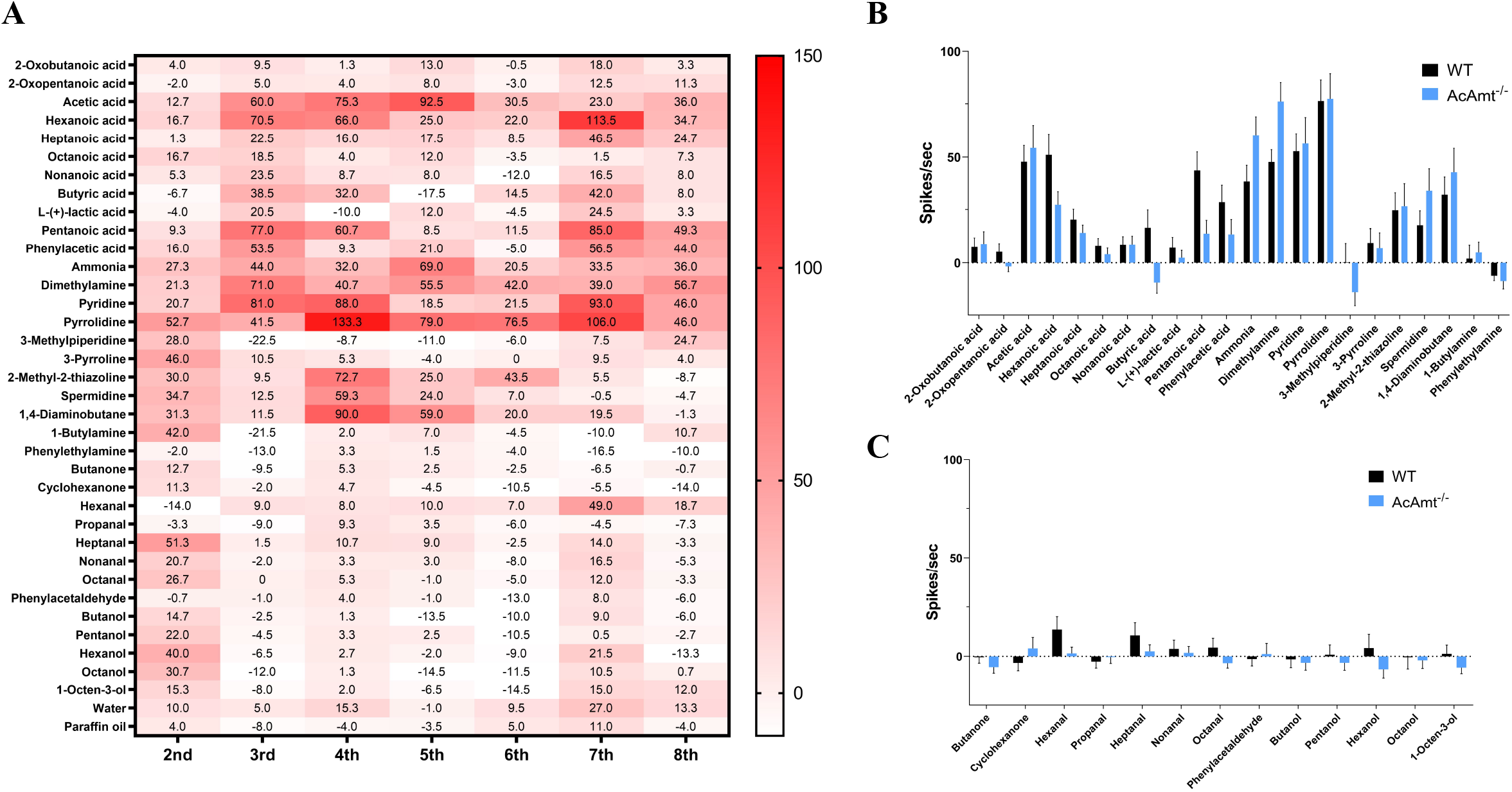
(**A**) Single-sensillum recording on wild-type female coeloconic sensilla. The heatmap showing mean responses to odorants (y-axis) in coeloconic sensilla on 2^nd^-8^th^ flagellomeres (x-axis; N=3-4 for each flagellomere); (**B**) Comparison of single-sensillum responses to amines and acids on total coeloconic sensilla between wild-type and *AcAmt*^-/-^ females (Multiple t-tests with Holm-Sidak method; N=2-4 for each from flagellomere 2^nd^-8^th^; N=23-25 in total); (**C**) Comparison of single-sensillum responses to ketones, aldehydes, and alcohols on total coeloconic sensilla between wild-type and *AcAmt*^-/-^ females (Multiple t-tests using Holm-Sidak method; N=2-4 for each from flagellomere 2^nd^-8^th^; N=23-25 in total). Error bars = Standard error of the mean.

Transcuticular EAG studies were also used to examine peripheral dose-dependent responses to ammonia at the whole-appendage level. Inasmuch as wild-type EAG responses to ammonia displayed both depolarization (downward) and hyperpolarization (upward) deflections relative to baseline (**Figure 4A**), these data were analyzed across both components. Once again, no significant differences were observed in dose-dependent antennal responses to ammonia (**Figure 4B&4C**) nor in response to the positive controls 1-octen-3-ol (**Figure 4D**) and butylamine (**Figure 4E**) which display robust dose-dependent depolarizations in both mutant and wild-type mosquitoes. Together, these data suggest ammonia responses across the antennae are not altered in *AcAmt*^-/-^ females. We also examined the role of *AcAmt* in peripheral responses to ammonia on the mosquito labella where it is also highly expressed (Pitts et al. 2014a; Ye et al. 2020) using ELG recording preparations. As was the case for the antennae, these studies demonstrated that both wild-type and *AcAmt*^-/-^ female labella display dose-dependent responses to ammonia with no significant differences (**Figure 4F**).

**Figure 4.**
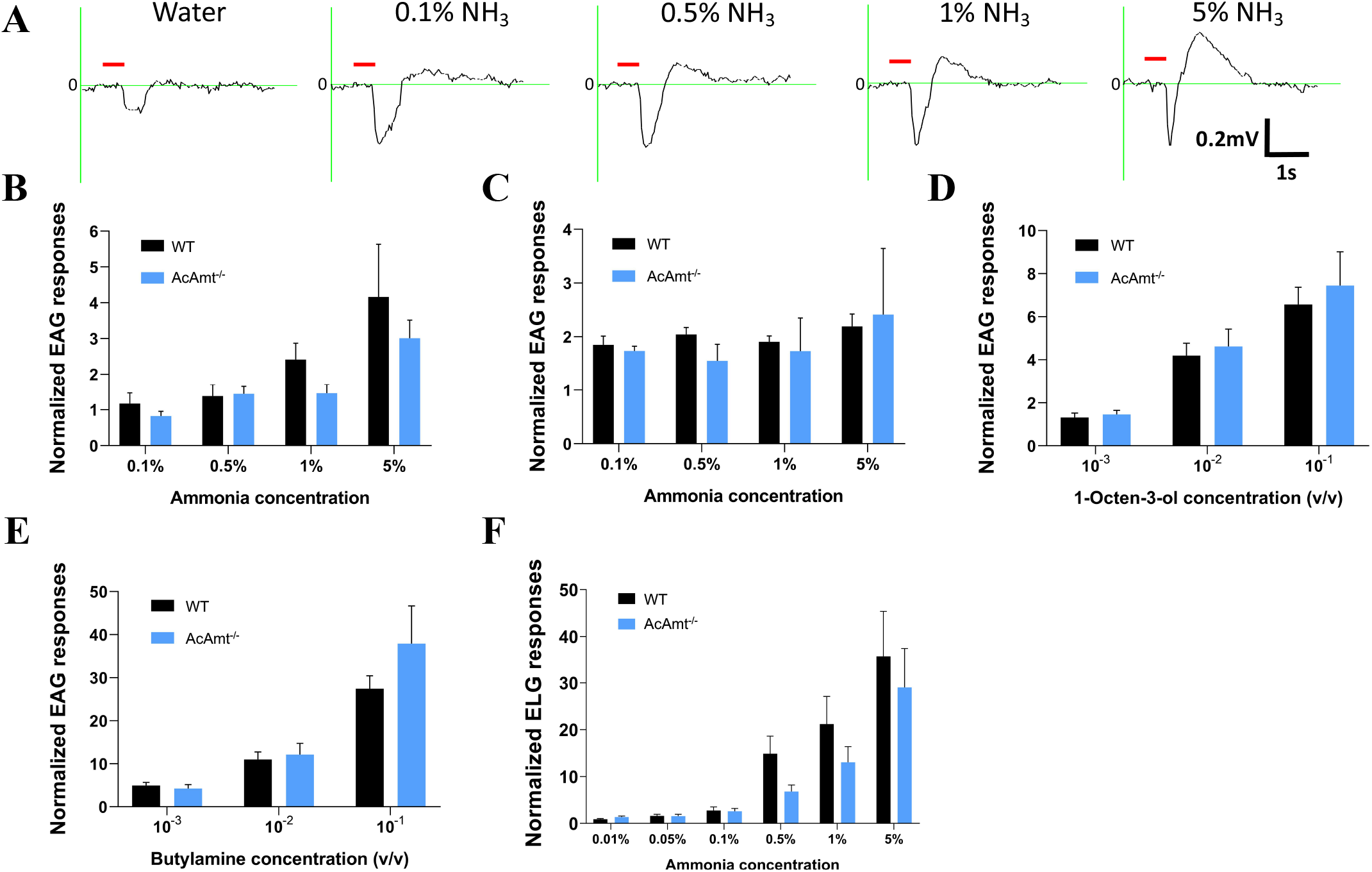
(**A**) Representative EAG responses of the wild-type to water and ammonia; Red bar indicates the duration of stimulations (0.5s). (**B**) Upward EAG responses to ammonia at different concentrations (N=6 for each concentration); (**C**) Downward EAG responses to ammonia at different concentrations (N=6 for each concentration); (**D**) EAG responses to 1-octen-3-ol at different concentrations (N=6 for each concentration); (**E**) EAG responses to butylamine at different concentrations (N=6 for each concentration); (**F**) ELG responses to ammonia at different concentrations (N=8-15 for each concentration). Multiple t-tests using Holm-Sidak method suggest no significant differences (P > 0.05) between the wild-type and *AcAmt*^-/-^. Error bars = Standard error of the mean.

### Reproductive deficits in *AcAmt* null mutants

In contrast to the absence of mutant olfactory phenotypes in response to ammonia, *AcAmt*^-/-^ mutants displayed a broad range of deficits associated with reproductive fitness and fecundity that resulted in a striking difficulty to propagate the *AcAmt*^-/-^ mutant line.

To assess this issue, we utilized a simple group mating bioassay (**Figure 5A**) to quantify female insemination rates (**Figure 5B**) which uncovered significant mating deficits in *AcAmt*^-/-^ mutants compared with the wild-type and *AcAmt*^+/-^ heterozygotes (**Figure 5C**). Importantly, this phenotype is not sex-specific as these mating deficits persist when pairing either female or male *AcAmt*^-/-^ mosquitoes with wild-type counterparts (**Figure 5C**). In order to investigate whether these phenotypes derived from shared or sex-independent mechanisms, we first examined male-specific processes such as sperm mobility. Here, *AcAmt*^+/-^ males, which produce both mutant and wild-type spermatozoa, were crossed with wild-type females thereby allowing the wild-type sperm to compete with mutant sperm throughout reproduction which is a multi-step process comprising insemination (i.e., the delivery of sperm to the female spermatheca) as well as subsequent sperm activation and oocyte fertilization. In this context, we quantified the number of heterozygous versus wild-type larvae distinguished by means of DsRed-derived fluorescence. In these studies, the consistent ratios of larval progeny showed there is no significant difference between the wild-type and *AcAmt* mutant sperm (**Figure 5D**). This suggests that the *AcAmt*^-/-^ mating deficits may be due to a reduction of the frequency of successful copulation, which raises the potential of broader deficits in overall metabolism that in turn impact general activity levels.

**Figure 5.**
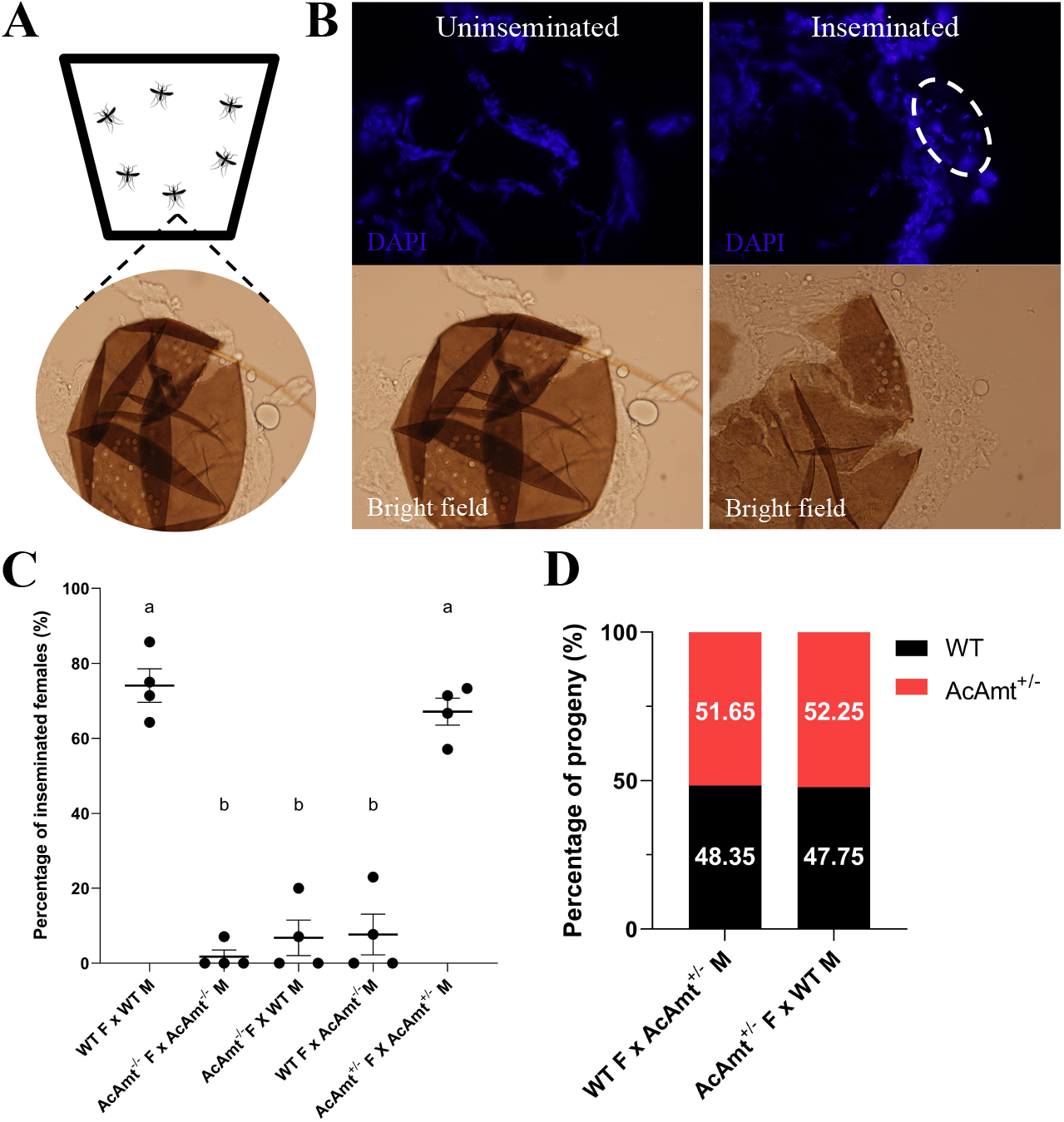
(**A**) Schematics of the mating bioassay. The females and males were allowed to mate in a bucket for 5 days before the female spermathecae were dissected; (**B**) Representation of inseminated and un-inseminated spermathecae stained with DAPI. The sperm heads are circled by dashed line; (**C**) Insemination rate of females in different mating pairs (F: females; M: males). Mean values with different grouping letters were significantly different (N=4; One-way ANOVA; P < 0.05); (**D**) Progeny ratio (wild-type versus *AcAmt*^+/-^) from two mating pairs to test sperm competency. Chi-square test suggests the ratio is not significantly different from 50% versus 50% (N=3-4; P > 0.05). Error bars = Standard error of the mean.

To assess activity profiles, individual male and female adult mosquitoes were digitally recorded in the scotophase between ZT12 and ZT17, which encompasses the peak period for Anopheline mating (Charlwood and Jones 1980; Howell and Knols 2009), and subsequently analyzed across several activity/mobility parameters, including distance travelled, the proportion of time spent near sugar water, the proportion of time spent moving, and the sum of clockwise and counterclockwise turns. Across the entire trial, both wild-type and *AcAmt*^-/-^ mutant females displayed a burst of activity within the first hour of the scotophase, followed by a prolonged period of relative quiescence (**Figure 6A**). Although the mean distance travelled over the full duration of the trial was relatively lower in males than females, a similar trend of activity and quiescence was also observed in both wild-type and *AcAmt*^-/-^ mutant males (**Figure 6B**). Furthermore, an analysis of all the activity/mobility parameters examined across the full duration of the bioassay failed to indicate any significant differences between wild-type or *AcAmt*^-/-^ mutant genotypes for either female or male adult mosquitoes (**Figure 6C-6J**). That said, these cumulative data largely reflect the prolonged period of inactivity, resulting in mean values that tend to converge the longer the mosquitoes remain inactive (**Figure 6A&6B**).

**Figure 6.**
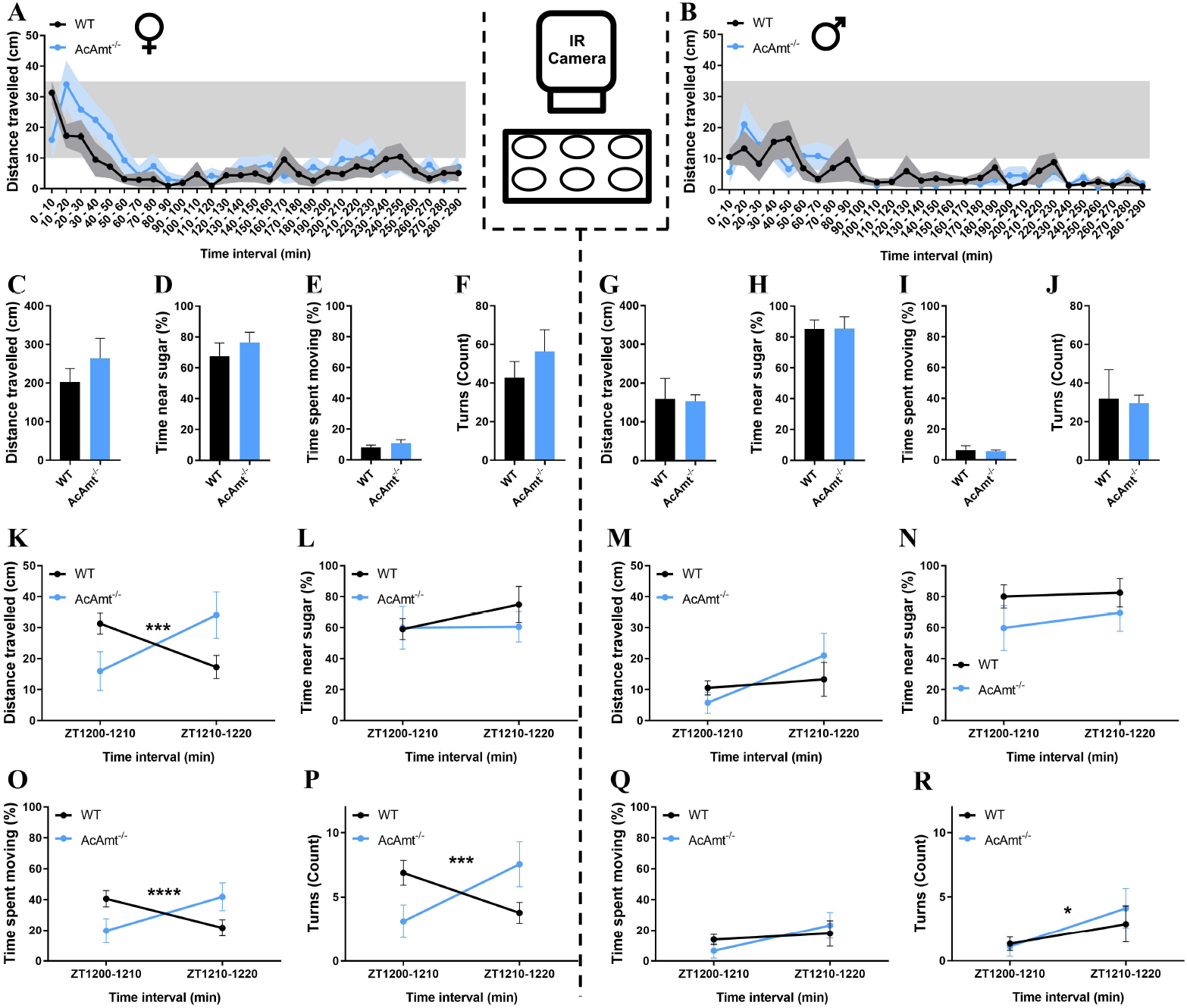
(**A-B**) Video recordings of individual mosquitoes showing distance travelled during the activity bioassay for wild-type and *AcAmt*^-/-^ mutant females (**A**) and males (**B**) organized into 10-min bins. Sex-specific data are separated by dashed lines. (**C-J**) Mobility parameters, including mean distance travelled (cm), time spent near the sugar water (%), time spent moving (%), and turning frequency (count) for females (**C-F**) and males (**G-J**) over the full duration of the bioassay (N=9; t-test with Welch’s correction); (**K-R**) Mobility parameters for females (**K-L**, **O-P**) and males (**M-N**, **Q-R**) over the first 20 min of the dark cycle organized into 10-min bins (N=9; Two-way repeated measures ANOVA; * < 0.05. *** = 0.0001. **** < 0.0001). Error bars = Standard error of the mean.

Inasmuch as the majority of mating in *An. coluzzii* occurs proximate to the dusk transition at start of the scotophase (Charlwood and Jones 1980; Howell and Knols 2009), we looked for more nuanced differences within this initial window. Here, wild-type females appeared to be more active within the first 10 min of the scotophase, while *AcAmt*^-/-^ mutant females manifested a modest latency in movement, which subsequently was higher than the wild-type females (**Figure 6A**). To address this more formally, we statistically analyzed activity levels within two discrete 10-min intervals that together represent the initial 20 min of the dark component of the light:dark cycle (ZT1200-1220). In this interval, while female mosquitoes showed no significant difference in the time spent near the sugar water, a significant interaction effect was observed between genotype and time with respect to distance moved (F(1, 16) = 26.28, P = 0.0001), the proportion of time spent moving (F(1, 16) = 27.99, P < 0.0001), and turning frequency (F(1, 16) = 24.59, P = 0.0001) (**Figure 6K-6L&6O-6P**). In males, apart from a modest but nevertheless significant ZT-dependent effect on turning frequency, in which both wild-type and *AcAmt*^-/-^ mutants turned more frequently in the ZT1210-ZT1220 interval than in ZT1200-ZT1210, no differences were observed (**Figure 6M-6N&6Q-6R**). Taken together, these results suggest that there are significant differences in activity levels between wild-type and *AcAmt*^-/-^ mutant females that correspond to the onset of the dark cycle and the peak period of mating (Charlwood and Jones 1980; Howell and Knols 2009). Specifically, *AcAmt*^-/-^ mutant females experience a delay in activity compared with their wild-type counterparts, which are most active at the onset of the dark cycle; this may contribute to mating deficiencies during this critical time window by desynchronization of peak activity between the sexes.

### Eclosion phenotypes

In addition to mating phenotypes, we also observed an interesting developmental deficit characterized by a significantly higher level of pharate mortality during eclosion of pupae to adults in *AcAmt*^-/-^ mutants compared with wild-type individuals raised under identical conditions and larval density levels (**Figure 7A&7B**). This phenotype does not appear to have a gender bias as approximately equal ratios of male and female *AcAmt*^-/-^ mosquitoes are represented in the reduced numbers of adults that nevertheless survive. Furthermore, *AcAmt*^-/-^ mosquitoes displayed the same pupation rate (**Figure 7C**) and general development timing as their wild-type and *AcAmt*^+/-^ counterparts, supporting the view that *AcAmt* mutations do not significantly influence larval or pupal stage development. Instead, these data suggest that post-eclosion reduction in viable *AcAmt*^-/-^ adults results exclusively from the failure of pharate adult *AcAmt*^-/-^ mutants to successfully eclose and fully emerge from their pupal cases. Taken together with the broad mating deficits of *AcAmt*^-/-^ mutants, these phenotypes raise the possibility that these mutants have an inability to effectively excrete or otherwise manage metabolic ammonia during these two intensively active processes resulting in toxic levels of ammonia that ultimately impact these critical behaviors.

**Figure 7.**
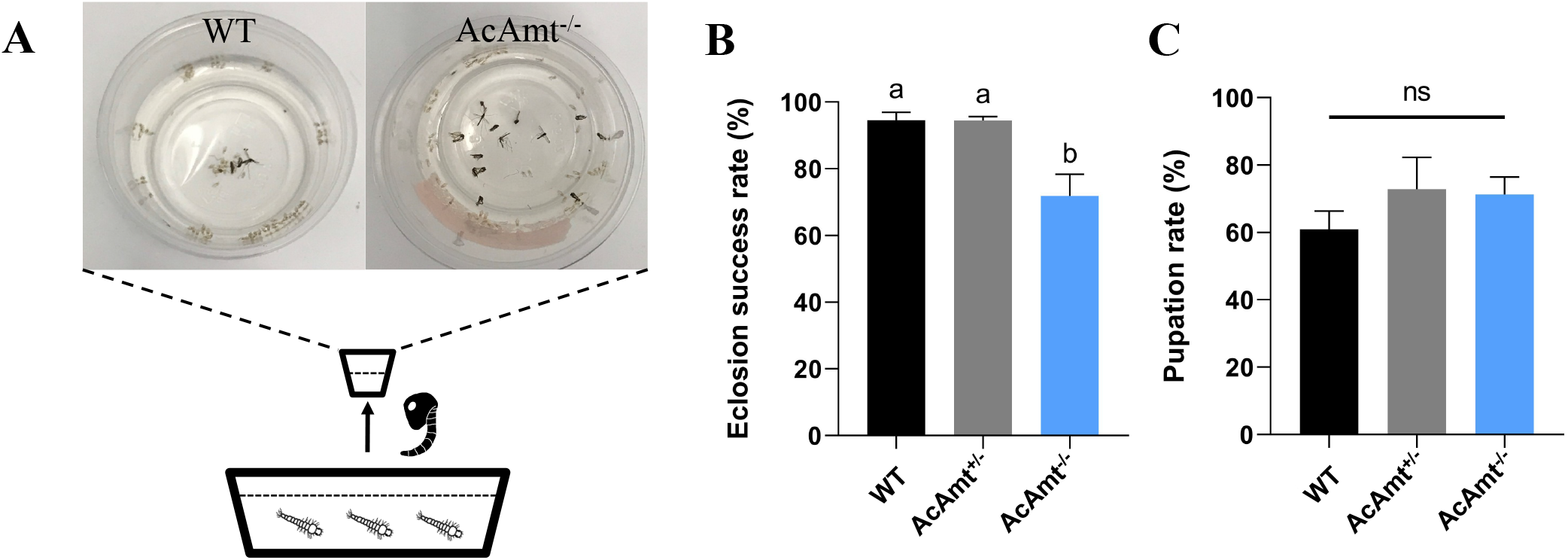
(**A**) Schematics of larval rearing in pan. Pupae were consequently placed in cups to examine eclosion rate. A representative image showing a higher mortality in *AcAmt*^-/-^ during eclosion; (**B**) Eclosion success rate. Mean values with different grouping letters were significantly different (N=5; One-way ANOVA; P < 0.05); (**C**) Pupation rate. One-way ANOVA suggests no significant differences among the three groups (N=5; P > 0.05). Error bars = Standard error of the mean.

### Elevated ammonia levels in *AcAmt* null mutants

RNAi-mediated silencing of *AeAmt1* has been shown to induce elevation of ammonia levels in the larval hemolymph of *Ae. aegypti* (Chasiotis et al. 2016). In order to assess this possibility in our *AcAmt*^-/-^ mutants, we used a simple colorimetric reagent to enzymatically measure whole-body ammonia levels in mating-stage adults and late-stage pupae (**Figure 8A**). These quantitative data indicate that while there was no alteration in ammonia levels for adult males regardless of genotype, mating-stage *AcAmt*^-/-^ females exhibited significantly higher levels of ammonia than wild-type females or *AcAmt*^+/-^ heterozygotes that displayed intermediate levels of ammonia (**Figure 8B**). Similarly, significant increases in ammonia levels were detected in unsexed late-stage *AcAmt*^-/-^ pupae relative to wild-type or *AcAmt*^+/-^ heterozygote counterparts (**Figure 8C**).

**Figure 8.**
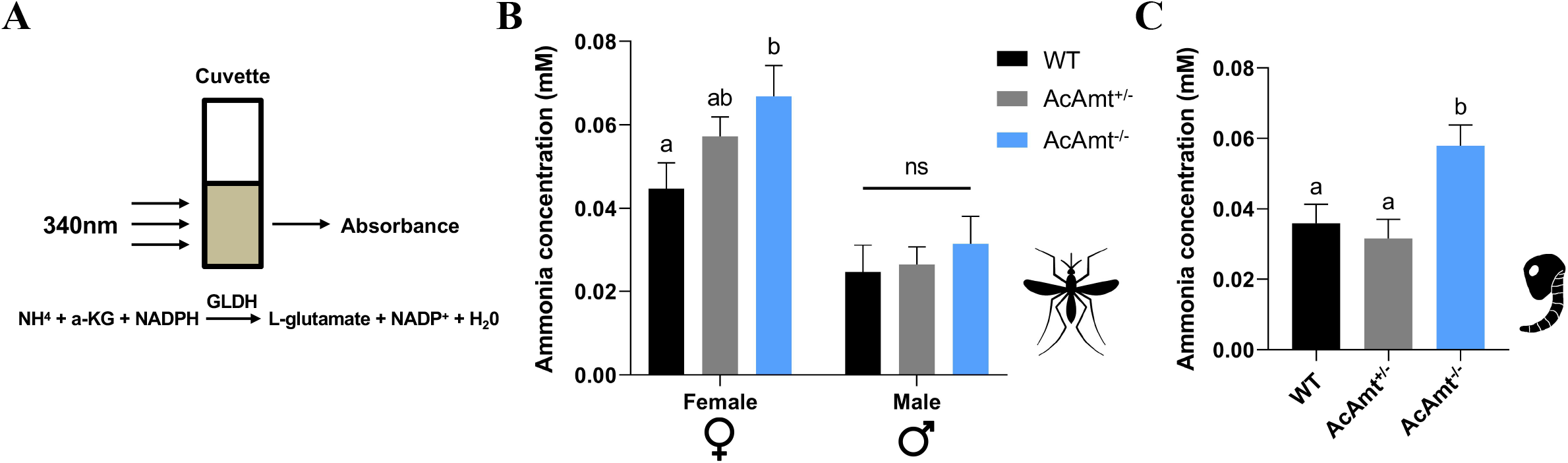
(**A**) Ammonia reacts with α-ketoglutarate (α-KG) and reduced nicotinamide adenine dinucleotide phosphate (NADPH) to form L-glutamate and NADP in a reaction catalyzed by glutamate dehydrogenase (GLDH) {L-glutamate: NAD(P) + oxidoreductase (deaminating), EC 1.4.1.3}, which is followed by a reduction of absorbance at 340nm; (**B**) Ammonia concentration in mosquito adults. Mean values with different grouping letters were significantly different (N=10-18; One-way ANOVA; P < 0.05); (**C**) Ammonia concentration in mosquito pupae. Mean values with different grouping letters were significantly different (N=12; One-way ANOVA; P < 0.05). Error bars = Standard error of the mean.

### Sugar feeding and carbohydrate levels in *AcAmt* mutants

The mating/eclosion phenotypes may also be the result of a potential defect in energy content and/or sugar feeding that play an essential role in mosquito mating and other behaviors (Gary et al. 2009). To examine this, we used a modified capillary feeder (CAFE) bioassay (**Figure 9A**) to measure adult sugar feeding during the same ZT12-ZT18 interval when Anopheline mating is most likely to occur (Howell and Knols 2009). In these studies, only *AcAmt*^-/-^ males exhibited a significantly lower sugar consumption than the wild-type and *AcAmt*^+/-^ males (**Figure 9B**). Water-only CAFE controls were also conducted, which demonstrated that the male-specific defect is restricted to sugar feeding (**Figure 9C**).

**Figure 9.**
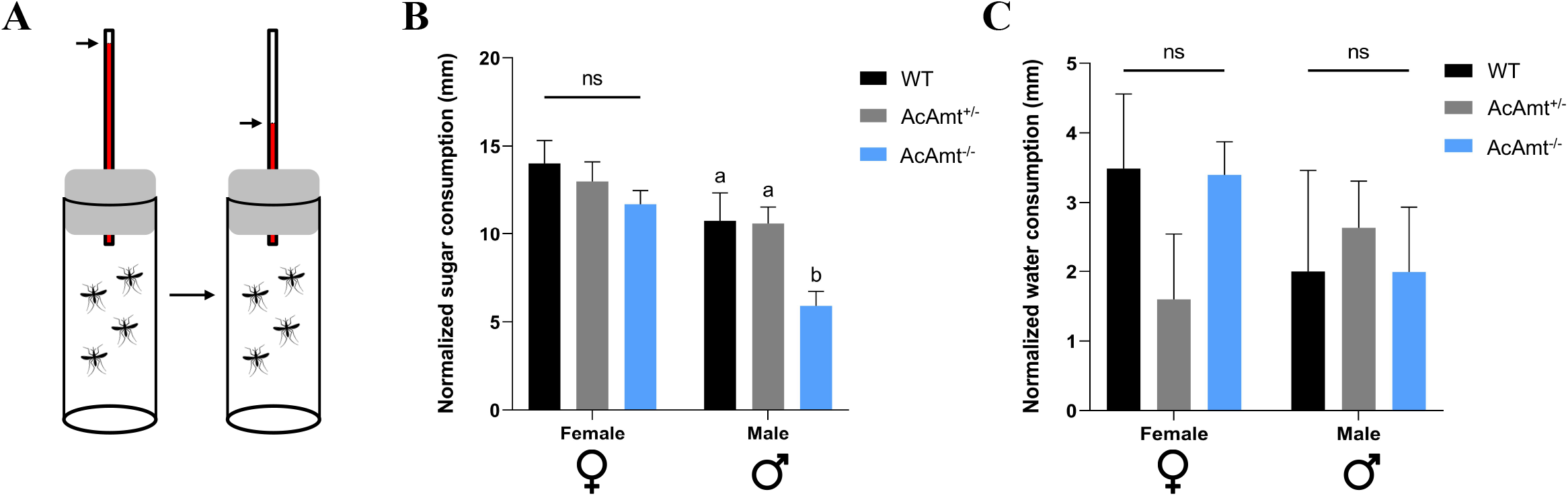
(**A**) Schematics of modified CAFE bioassay where the consumption was quantified by sugar level reduction marked on the capillary; (**B**) Sugar feeding ability in mosquito adults. Mean values with different grouping letters were significantly different (N=6-8; One-way ANOVA; P < 0.05); (**C**) Water consumption controls in mosquito adults. One-way ANOVA suggests no significant differences among the three groups (N=6-8). Error bars = Standard error of the mean.

To further examine the potential impact of sugar feeding deficits on adult mating and pupal eclosion, respectively, we collected adults at ZT11 just before the onset of mating and late-stage pupae 12h before eclosion and used the phenol-sulfuric acid method (Ahmed 2013; Ellison et al. 2015) to assess whole-body carbohydrate levels across wild-type and *AcAmt* mutant genotypes. Once again, while there were no significant differences across adult male genotypes, *AcAmt*^-/-^ females exhibited significantly lower total carbohydrate content than wild-type or the intermediate levels seen in *AcAmt*^+/-^ heterozygotes (**Figure 10A&10B**). In order to control for larger individuals artifactually accounting for these higher carbohydrate contents, mosquitoes were sampled and weighed prior to homogenization. Correspondingly, this analysis revealed that both *AcAmt*^-/-^ and *AcAmt*^+/-^ females weighed significantly less than wild-type females (**Figure 10C**), which suggests their lower carbohydrate contents may, in part, reflect this physical characteristic.

**Figure 10.**
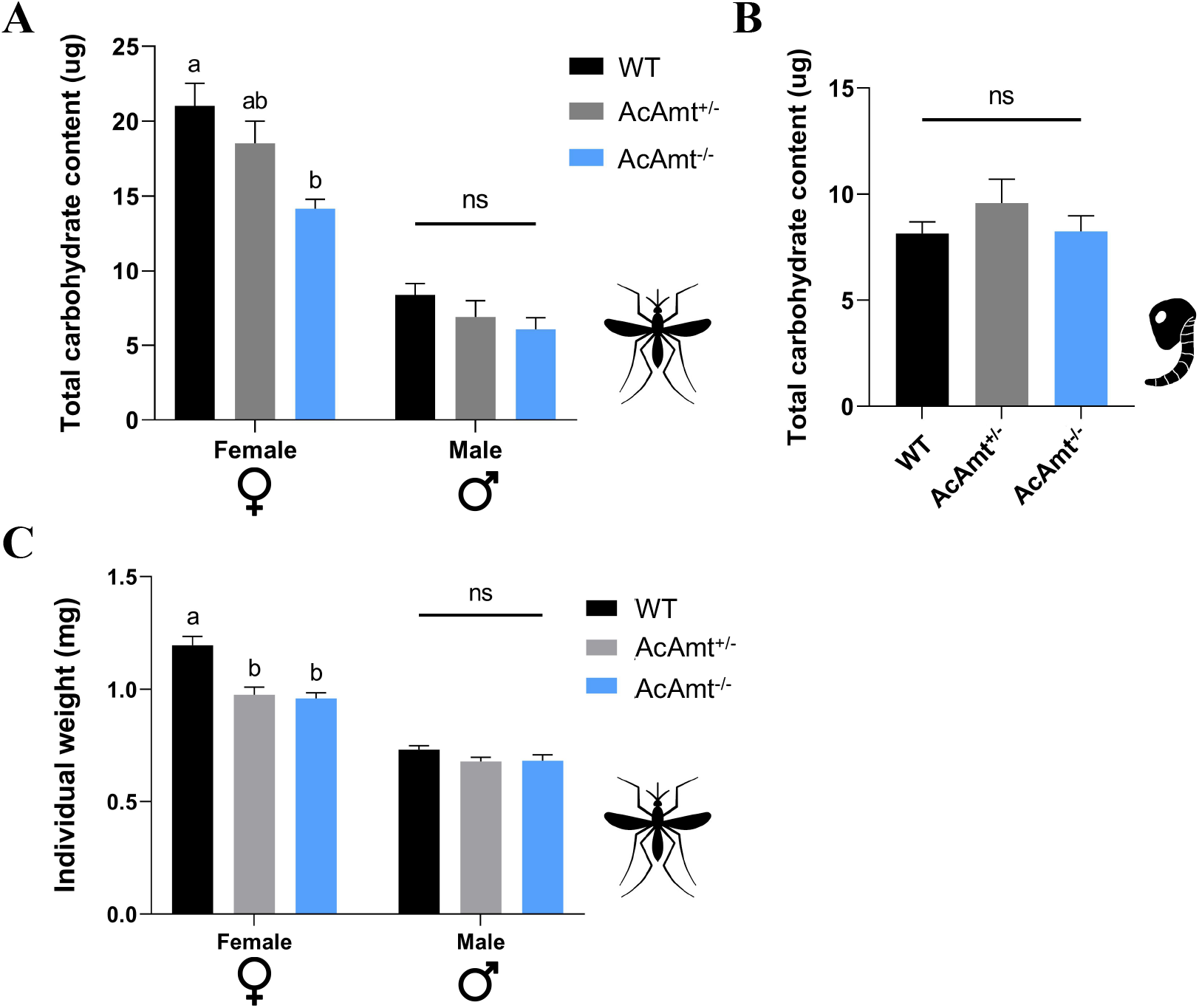
(**A**) Total carbohydrate content in mosquito adults. Mean values with different grouping letters were significantly different (N=8; One-way ANOVA; P < 0.05); (**B**) Total carbohydrate content in mosquito pupae. Mean values with different grouping letters were significantly different (N=6; One-way ANOVA; P < 0.05); (**C**) Individual mosquito adult weights. Mean values with different grouping letters were significantly different (N=10-15; One-way ANOVA; P < 0.05). Error bars = Standard error of the mean.

## Discussion

In *Drosophila, DmAmt* null mutants demonstrated a dramatic reduction of ac1 sensilla responses to ammonia where *DmAmt* is expressed in auxiliary cells and hypothesized to be involved in ammonium clearance (Menuz et al. 2014), while no such phenotype was observed in the labella where *DmAmt* is exclusively neuronal (Delventhal et al. 2017). We now report a comprehensive investigation in the malaria vector mosquito *An. coluzzii* of *AcAmt* null mutant olfactory responses to ammonia. This analysis encompasses both antennal grooved pegs and coeloconic sensilla, where *AcAmt* is primarily expressed in auxiliary cells, as well as the labella where *AcAmt* was observed in olfactory and non-olfactory neurons (Ye et al. 2020). In contrast to *Drosophila*, no significant reduction of peripheral neuron sensitivity to ammonia was found in either antennae or labella of *AcAmt*^-/-^ mutants. It is noteworthy that, in addition to *AcAmt*, another ammonium transporter, *Rh50*, is highly expressed on the mosquito antennae (Pitts et al. 2014a), which in light of these data is to likely play a complementary role in the ammonia-sensing and management pathways. This is consistent with a previous study in *Drosophila*, in which ammonia responses in ac3 and ac4 sensilla on female antennae where *DmRh50* is expressed were not impacted by the *DmAmt* mutation (Menuz et al. 2014).

While the receptors and other components underlying ammonia-sensing mechanisms in the mosquito olfactory system remain unknown, attraction to ammonia plays a significant role in host-seeking behaviors by *Anopheles* females (Braks et al. 2001; Smallegange et al. 2005). This makes it likely that sensitivity to ammonia is sufficiently essential in anautogenous mosquitoes to drive the evolution of parallel and complementary ammonia sensitivity processes. In light of the lack of *AcAmt*^-/-^ deficits in *An. coluzzii* olfaction, comprehensive localization and characterization of the ammonium transporter *Rh50* will be critically informative. Indeed, it is reasonable to speculate that significant impairment of olfactory responses to ammonia might require mutations of both *AcAmt* and *Rh50*.

In addition to the peripheral olfactory responses to ammonia, ammonium transporters have recently been shown to be involved in other essential functions in the biology of insects including male fertility in *Ae. aegypti* (Durant and Donini 2020) and larval muscle control in *Drosophila* (Lecompte et al. 2020). Here, CRISPR/Cas9-induced *AcAmt*^-/-^ mutations similarly uncover several potentially non-olfactory phenotypes in *An. coluzzii* that are likely to significantly reduce the overall fitness of these mutants. Even so, while significant fecundity deficits are reported here in *AcAmt*^-/-^ mutants and *AeAmt* RNAi treatments in *Ae. aegypti* (Durant and Donini 2020), these phenotypes are likely to result from fundamentally different mechanisms working synergistically. In *An. coluzzii*, the frequency of successful copulation (sperm delivery) is significantly reduced in both *AcAmt*^-/-^ females and males, while the decrease of fecundity in *Ae. aegypti* appears to be due to a significant reduction in viable spermatozoa (Durant and Donini 2020). Importantly, this latter phenotype is specifically not observed in *An. coluzzii* mating studies. Instead, data reported here suggest that the absence of *AcAmt* results in subtle but nevertheless significantly altered activity profiles during the circadian interval most associated with mating (Charlwood and Jones 1980; Howell and Knols 2009). Even more compelling is the *AcAmt*-dependent elevation of endogenous ammonia levels that may rise above physiologically toxic thresholds in mating-stage adults and late-stage pupae as the likely mechanism responsible for these mating as well as the eclosion deficits we report. This rationale aligns with increased ammonia levels and hemolymph acidification found in *Ae. aegypti* larvae treated with *AeAmt/AeRh50*-targeted RNAi (Chasiotis et al. 2016; Durant et al. 2017; Durant and Donini 2018) and suggests that *AcAmt* is similarly involved in ammonia management/excretion systems in *Anopheles* mosquitoes. This is consistent with our recent hypothesis implicating *AcAmt* in neural toxicity and ammonia homeostasis (Ye et al. 2020).

During mating, both male and female mosquitoes monitor each other’s wing beat frequency to actively modulate these activities toward convergence (Gibson and Russell 2006; Cator et al. 2009; Robert 2009; Gibson et al. 2010). This auditory interaction between females and males has been suggested to serve an important role in conspecific mating recognition and, in that context, directly contributes to mosquito reproductive fitness (Cator et al. 2009; Robert 2009). This has indeed been shown to contribute to the reproductive isolation between “M” and “S” forms of *An. gambiae* now recognized as distinct species (Coetzee et al. 2013), which utilize wing beat frequency to recognize potential mates within their own molecular form/species (Pennetier et al. 2010). Notably, this mating interaction requires not only auditory interactions, but also the coordination of wing movement to match the frequencies of corresponding partners (Robert 2009). While uncharacterized in mosquitoes, *Drosophila* leg and wing muscles are innervated with glutamatergic neurons, and, not surprisingly, the malfunction of these neurons impairs fly movement (Sadaf et al. 2015; Gowda et al. 2018). Inasmuch as *Anopheles* mosquitoes rely on muscle coordination to achieve a matching of wing-beat frequencies between females and males for mating recognition (Pennetier et al. 2010), the absence of *AcAmt* function may impact neuronal function to impair muscle control and the auditory/wing beat frequency convergence required during mating. With regard to the eclosion deficits displayed by *AcAmt* mutants, it is reasonable to conclude that successful emergence from the pupal case requires similarly substantial muscular coordination and effort such that failure to physiologically manage ammonia/acid levels could well be lethal.

It appears likely that multiple complementary systems exist in mosquitoes to ensure ammonia detection, which is critical for host seeking and reproduction. Similarly, it seems likely that Amts, Rh50s, as well as other cryptic ammonium transporters are involved in distinct functional pathways where they play essential roles in supporting locomotion and behavior. Taken together with our recent *AcAmt* localization study (Ye et al. 2020), the CRISPR/Cas9-mediated genome-editing studies reported here suggest that *AcAmt* is functional across a variety of systems that involve olfaction, reproduction, and ammonia metabolism. Whereas further integrative studies on different ammonium transporter genes will doubtlessly reveal more detail regarding these functions, the broad footprint of *AcAmt* activity, especially insofar as its impact on mosquito fecundity, supports its role as an important target for the development of novel vector-control strategies.

---

**Supplementary Table 1.** The oligonucleotide primers used in this study. (**A**) sgRNA oligos targeting *Exon 1* of *AcAmt*; (**B**) sgRNA oligos targeting *Exon 4* of *AcAmt;* (**C**) Primers amplifying the homologous arm extending from the DSB site in *Exon 1* which was inserted into the *AarI* site of the homologous template; (**D**) Primers amplifying the homologous arm extending from the DSB site in *Exon 4* which was inserted into the *SapI* site of the homologous template; (**E**) Primers used in the PCR confirmation of *AcAmt* mutagenesis.

## Acknowledgements

We thank Zhen Li for mosquito rearing and all members of the Zwiebel lab for critical suggestions, as well as Drs. Julian Hillyer, Maulik Patel, Wenbiao Chen, and Patrick Abbot (Vanderbilt University) for valuable advice throughout the course of this work. We also thank Dr. Samuel Ochieng for technical support in conducting ELGs, Dr. Willi Honegger for comments on the manuscript and Dr. AM McAinsh for scientific copy-editing. This work was conducted with the support of Vanderbilt University and funded by the National Institutes of Health (NIAID, R21-113960) to RJP and LJZ.

## Declarations

### Funding

This work was conducted with the support of Vanderbilt University Endowment Funds and by a grant from the National Institutes of Health (AI113960) to LJZ.

### Conflicts of interest

The authors declare that they have no competing interests.

### Availability of data and material

All data generated or analyzed during this study are included in this published article and its supplementary information files.

### Author contributions

Conceived experiments: ZY, FL, STF, RJP and LJZ; Performed research: ZY, FL, and STF; Analyzed data: ZY, FL, STF, and AB; Wrote the paper: ZY, FL, STF, AB, RJP, and LJZ. Approved the final manuscript: ZY, FL, STF, AB, RJP, and LJZ.

## Notes

### Competing Interest Statement

The authors have declared no competing interest.

